# A Cyclic Arginine Adduct Eclipses Carboxymethylation as the Primary Glyoxal-Derived Advanced Glycation End-Product

**DOI:** 10.1101/2025.05.27.656459

**Authors:** Morgan E. Brutus, Colby S. Girard, Jeremiah W. Jacob-Dolan, Rebecca A. Scheck

## Abstract

Glyoxal (GO) is a highly reactive 1,2 dialdehyde implicated in the formation of a set of disease-associated post-translational modifications (PTMs) known as advanced glycation end products (AGEs). While GO has been widely reported to modify lysine to form the highly studied AGE carboxymethyllysine (CML), here we demonstrate that GO alone leads to highly chemoselective arginine glycation, yielding a stable glyoxal-derived hydroimidazolidine (GH-DH) product. This near-exclusively formed AGE has the same mass change as carboxymethylarginine (CMA), which implies that it may have been overlooked or misattributed in prior studies. In contrast, lysine modification by GO is highly dependent on the presence of hydride-based reducing agents, suggesting that prior reports may have artifactually generated CML through pervasive use of reductive amination protocols during sample preparation. These findings challenge the standing assumptions about the landscape of GO-derived glycation, emphasizing the importance of carefully considering the impact of experimental conditions in glycation studies. By redefining the complement of GO-derived AGEs, this study provides essential new information that is greatly needed for uncovering their biology, creating new tools for their study, and discovering therapies that ameliorate or mitigate their glycation-related damage.

---

Glyoxal (GO) is the smallest of the α-oxoaldehydes, a 1,2 dialdehyde, that forms through multiple pathways in biological systems, including lipid peroxidation and glucose autoxidation^[1– 3]^. Typical serum concentrations of glyoxal are estimated to be around 220 nM, but can increase by more than threefold under metabolic dysregulation associated with diseases including diabetes, uremia, and renal disease.^[4–6]^ Given its elevated levels in disease and its enhanced electrophilicity, GO is known to react non-enzymatically with nucleophilic residues in proteins, predominately targeting lysine and arginine side chains in a process known as glycation^[7–9]^. Glycation produces a group of post-translational modifications (PTMs) known as advanced glycation end products (AGEs). In particular, GO (58.04 Da) is reported to produce at least four discrete AGEs, including carboxymethyllysine (CML, [M+58]), the glyoxal-derived hydroimidazolone (GH-1, [M+40]) and hydroimidazolidine (GH-DH, [M+58]) isomers, and carboxymethylarginine (CMA, [M+58]) (**Figure 1**)^[1,10,11]^.

**Figure 1.**
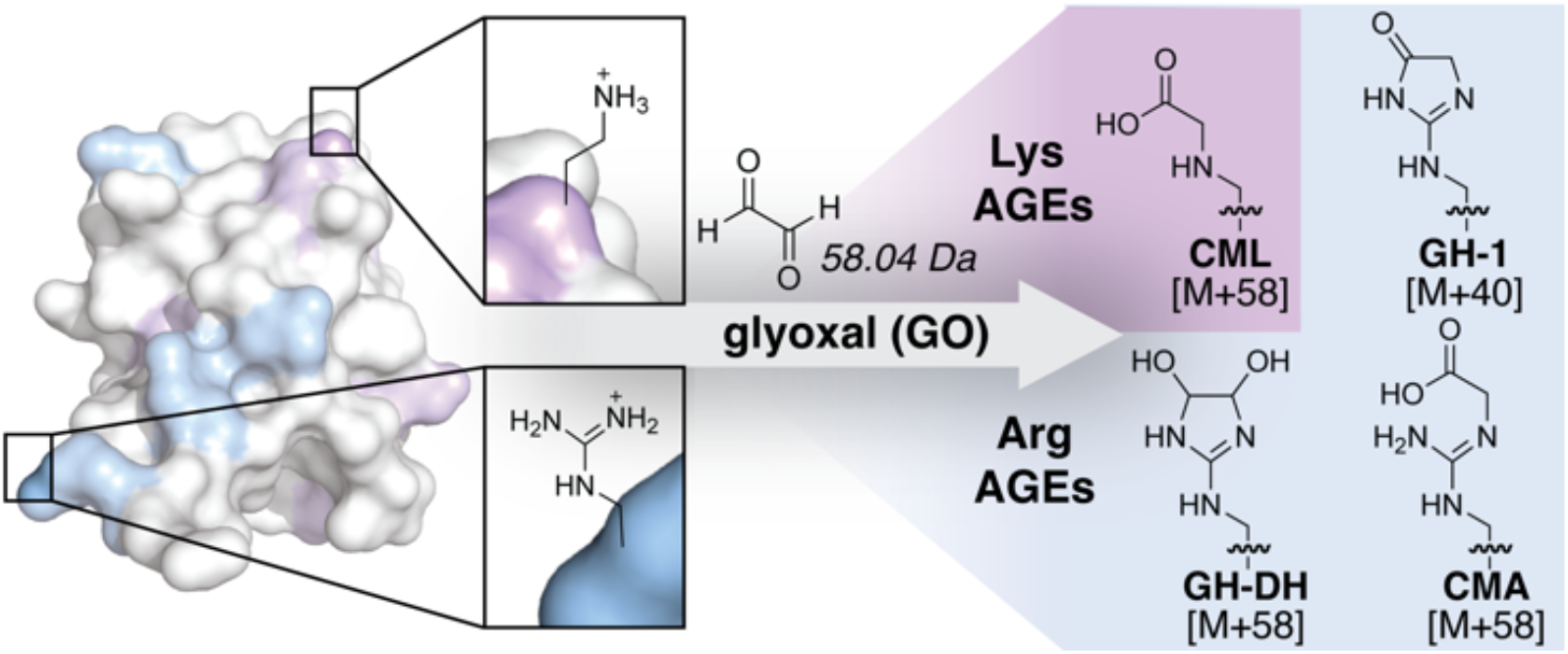
Previously reported glyoxal (GO)-derived advanced glycation end-products (AGEs).

Among these, carboxymethylation of Lys (CML formation) is particularly well-known, as it has been extensively studied as a biomarker for glycation-related protein damage^[12–15]^. CML is thought to be among the most abundant AGEs and has been implicated in numerous pathologies, including diabetes^[16]^, atherosclerosis^[17]^, chronic kidney disease^[18]^, and neurogenerative disorders^[19]^. CML accumulates in long-lived extracellular matrix proteins such as collagen, contributing to tissue stiffening and loss of function over time^[20,21]^. CML is known to trigger inflammatory and oxidative stress pathways that exacerbate disease progression through the receptor for advanced glycation end-products (RAGE)^[22,23]^. Moreover, CML has been identified in histones, where it may influence chromatin structure and gene expression^[24,25]^. Recently, CML was shown to disrupt proteostasis by triggering a proteotoxic response, impairing protein degradation, and affecting cellular functions such as microtubule dynamics and cell cycle regulation^[26]^.

Given the significance of CML as a biomarker for glycation-related damage and disease risk, and its potential as a driver for disease, it is critical to understand the specific metabolites that are responsible for its formation. However, relatively little is known about the propensity of GO to modify lysine (forming CML) over arginine (producing CMA, and the GH or GH-DH isomers (n. b. only GH-1 and GH-DH-1 are shown in **Figure 1**). In this study, we evaluate the ability of GO to modify Lys or Arg residues on peptides and proteins. Our findings demonstrate that GO predominantly targets Arg, forming a single, remarkably persistent dihydroyimidazolidine (GH- DH) product that endures over extended periods without appreciable conversion to other AGEs. Conversely, lysine modification by GO is highly dependent on the addition of hydride reducing agents, suggesting that commonly used protocols for studying CML may have produced it artifactually. Learning the major AGEs produced by each biologically relevant glycating agent is essential for uncovering their roles in disease and developing targeted strategies to ameliorate or mitigate their glycation-related damage.

To begin, we evaluated the reaction of GO with a hit peptide (Ac-LESRHYA, peptide **1**^**R**^) that we identified in a previous study^[27]^. Compared to the related α-oxoaldehyde methylglyoxal (MGO), we found that GO was far less reactive in modifying Arg residues despite the presence of two aldehydes. Only 22.6 ± 0.9% glycation was observed for peptide **1**^**R**^ (1 mM) treated with 1 mM GO for 24 hours, while 40.1 ± 0.8% glycation was observed by MGO under the same treatment conditions (**Figure S1**). To ensure that reduced glycation with GO was not due to major discrepancies in glycating agent concentrations, we used a derivatization assay, which revealed the commercial stocks of GO to even more concentrated than those of MGO (**Figure S2**). We therefore suspect that the dampened glycation reactivity of GO arises from its tendency to form a dihydrate in aqueous solutions, which may slow initial addition steps. Moreover, while MGO produced a range of AGEs on peptide **1**^**R**^, GO-derived glycation produced just a single product with a [M+58] mass change, even when higher concentrations of GO (5 mM) were used. This adduct (78.8 ± 8.3%) appears as a single peak that elutes with a similar retention time to the parent peptide (**Figure 2a,b**). While this could be either CMA or GH-DH based on mass change alone, we suspected it was GH-DH, as our previous mechanistic work with MGO revealed that the dihydroxyimidazolidine (MGH-DH) forms initially. Indeed, using a chemical derivatization assay that tests for 1,2 diols, along with 2D NMR, we confirmed that this product matched the structure of GH-DH, and is most consistent with the GH-DH-1 isomer (**Figures S3 and S4**).

**Figure 2.**
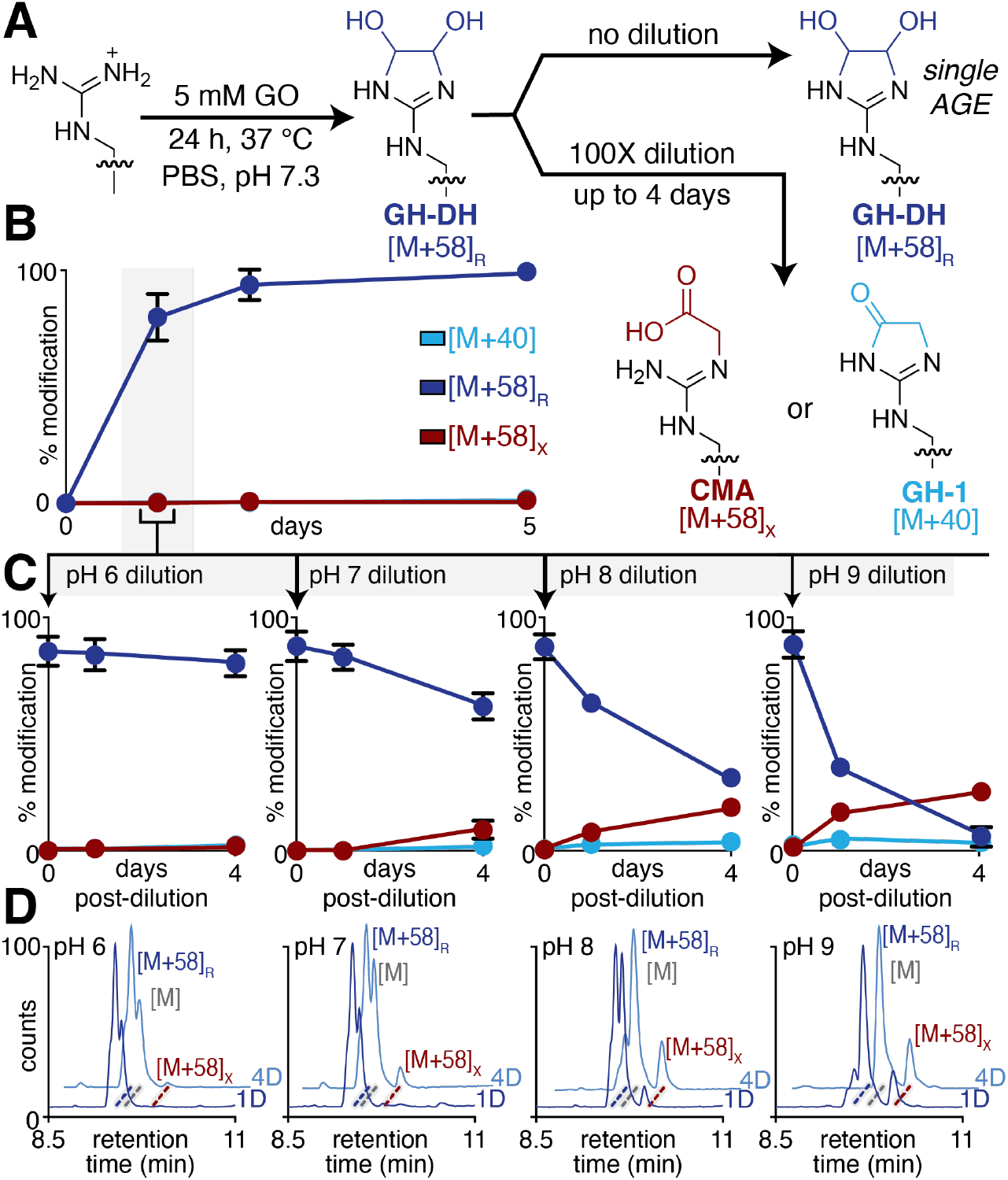
Evaluating Arg Glycation by GO. A) Reaction scheme for peptide glycation with GO. B) Time course data for 1 mM peptide **1**^**R**^ treated with 5 mM GO in PBS pH 7.3 for up to 5 days at 37 °C. LC-MS analysis along with structure determination by 2D NMR revealed that an [M+58]_R_ adduct corresponding to GH-DH is the only AGE formed. C) Dilution of the GO glycation reaction after 24 h, into PBS at pH ranging from 6-9 shows that GH-DH is persistent at neutral pH, with only small quantities of [M+58]_X_ or [M+40] adducts forming after 4 days of incubation at elevated pH, as shown in the quantified LC-MS time course data and D) representative base peak ion chromatograms.

Our previous mechanistic work also revealed that MGH-DH is a precursor to the hydroimidazolone, MGH-1, which can hydrolyze into carboxyethylarginine (CEA)^[27]^. Given the structural similarity between GO and MGO, we expected that a similar mechanistic pathway should be possible, where GH-DH is eliminated to form GH-1, which could subsequently hydrolyze resulting in the formation of carboxymethylarginine (CMA) (**Figure 2a**). To evaluate this possibility, we used a dilution protocol we previously developed to monitor the rearrangement of GH-DH into other AGEs^[27]^. Unexpectedly, GH-DH exhibited remarkable stability, persisting for up to four days at pH 7 (**Figure 2c,d**). Although there was a 25.3 ± 5.7% loss of GH-DH over the four days post-dilution, most was lost to reversal of glycation, regenerating unmodified peptide **1**^**R**^. Only a small, but appreciable amount (9.4 ± 3.3%) of a new, distinct [M+58]^X^ adduct that we suspected to be CMA was observed after four days post-dilution.

To further evaluate this behavior, we performed the same dilution protocol across a range of pH. We observed that GH-DH exhibited even greater temporal stability at pH 6, whereas substantial removal occurred when the reaction was diluted at pH 8 or 9. After four days, only 30.1 ± 2.0% of GH-DH remained at pH 8, and this decreased further to 5.7 ± 3.3% at pH 9. In both cases, some of the lost GH-DH converted to the [M+58]_X_ adduct, but a far greater portion reverted to unmodified peptide (37.1 ± 3.7% and 55.5 ± 6.5% at pH 8 and 9, respectively). Additionally, we observed a minor product with an [M+40] mass change, likely corresponding to the GH isomers (**Figure 2c,d**). These results suggest that unlike the highly abundant and persistent MGH isomers, GH isomers are significantly less stable.

Next, we evaluated the reaction of GO and MGO on a Lys-bearing version of our hit peptide, peptide **1**^**K**^ (Ac-LESKHA). Although GO and MGO are widely reported to form CML and CEL^[11,28–33]^, respectively, we did not observe any modification of peptide **1**^**K**^, even using 5 mM GO at 37 °C for 24 hours (**Figure 3**). This result contrasts with our findings for the Arg-containing peptide, where substantial modification was detected under identical conditions. To us, the lack of Lys modification suggests a difference in reactivity, as dicarbonyls like MGO and GO preferentially modify arginine, likely due to the bidentate binding interaction with the guanidinium group, which stabilizes cyclic adducts through a chelate effect^[7]^. In contrast, the Lys ε-amine, which lacks these interactions, is less reactive toward dicarbonyls under these conditions, especially as these small dicarbonyls lack an ?-hydroxyl group that is required for the Amadori rearrangement^[34]^.

**Figure 3.**
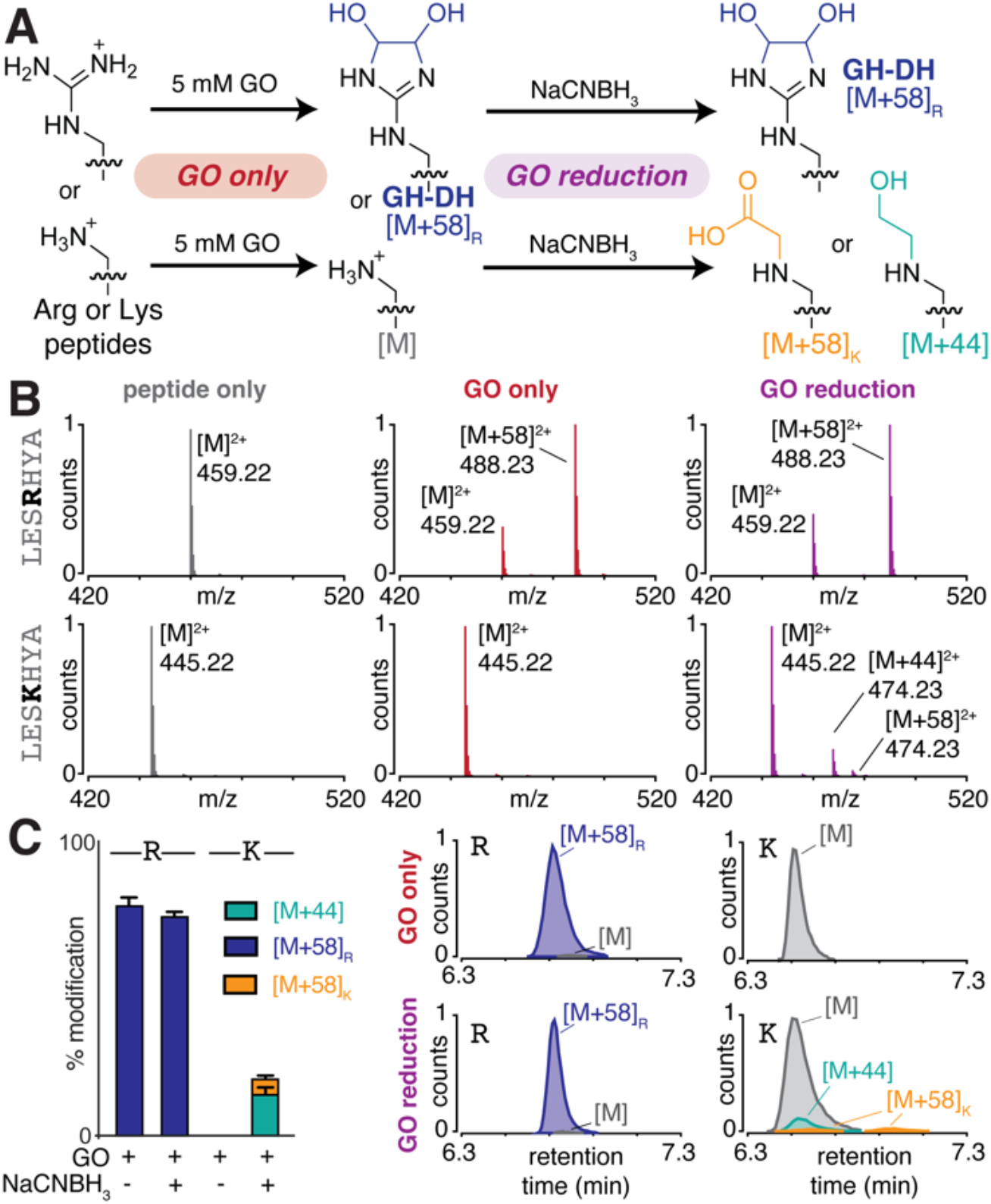
Evaluating Lys Glycation by GO. A) Reaction scheme for modification of Arg or Lys peptides by GO, both alone (GO only) and with subsequent addition of sodium cyanoborohydride (GO reduction). B) Representative LC-MS spectra for 1 mM peptide **1**^**R**^ or **1**^**K**^ treated with 5 mM GO in PBS pH 7.3 for 24 h at 37 °C. C) Quantified LC-MS data shows the distribution of all modified products under each conditions tested, along with representative total compound chromatograms (TCC). Together, these results clearly demonstrate that only peptide **1**^**R**^ is glycated by GO, while **1**^**K**^ is modified only under the GO reduction conditions.

Nonetheless, there is substantial literature precedence for GO as a primary source for CML formation^[11,35]^. To reconcile our observations with past reports, we carefully evaluated the methods used to obtain CML in prior studies. Crucially, we noted that every instance of reported GO-mediated CML formation involved a reduction step using hydride reducing agents^[11,24,36–40]^. To test whether reduction influenced the complement of GO modifications, we treated peptides **1**^**R**^ and **1**^**K**^ with sodium cyanoborohydride following treatment with 5 mM GO for 24 h at 37 °C (**Figure 3a**)^[41]^. Using this protocol, we found that the AGE distribution for peptide **1**^**R**^ remained similar before and after treatment with sodium cyanoborohydride. However, for peptide **1**^**K**^, modest levels of modification were now clearly discernable. The major product (15.0 ± 2.1%) had a mass change of [M+44], matching with the expected reductive amination product in which both the Lys-GO Schiff base is reduced along with the remaining aldehyde on GO (**Figure 3a-c**). Additionally, small amounts of an [M+58]_K_ Lys adduct were observed by LC-MS at two different retention times, one at 6.6 min and the other at 7.0 min, suggesting they are two different AGEs (**Figure 3c**). These findings indicate that Arg readily forms stable AGEs with GO under physiological conditions, whereas Lys modification requires reductive amination.

To evaluate if these results extended to protein substrates, we conducted a similar set of experiments using three different proteins: ubiquitin (4 Arg, 7 Lys), RNAse (4 Arg, 9 Lys), and lysozyme (11 Arg, 6 Lys). Each protein was treated with 5 mM GO at 37 °C for 24 h, with or without subsequent sodium cyanoborohydride treatment, and followed by LC-MS analysis (**Figure 4a**). When treated with GO only, the deconvoluted spectra reveal clear modified peaks, appearing in a pattern of single, double, or triple [M+58] additions detected across all proteins tested. Upon addition of sodium cyanoborohydride (GO reduction), the spectra indicate a higher degree of modification, displaying a mixture of [M+44] and [M+58] adducts. Subsequent proteolytic digestion and MS/MS analysis confirmed that GO exclusively modified Arg residues when no reducing agent was added. However, when sodium cyanoborohydride was introduced post-GO treatment, both Arg and Lys were modified (**Figure S5**).

**Figure 4.**
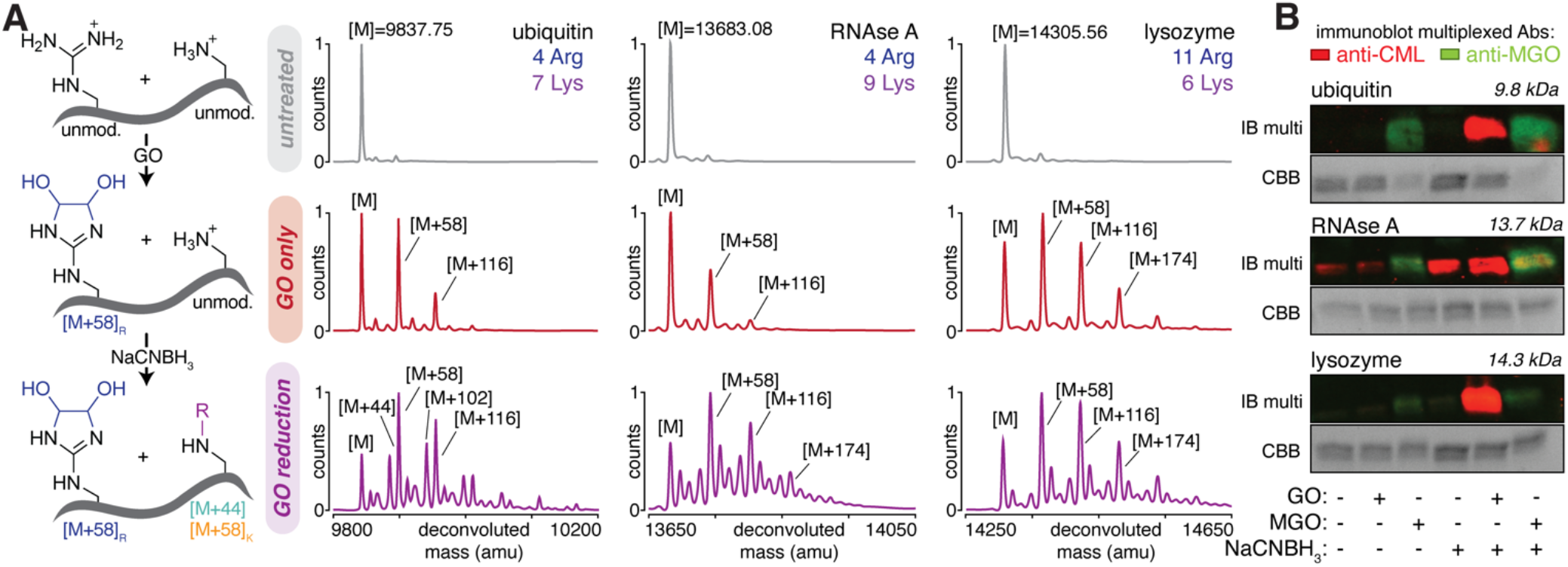
Assessing Protein Glycation by GO. A) Reaction scheme alongside representative deconvoluted intact protein mass spectra showing GO glycation using GO only or GO reduction conditions for ubiquitin, RNAse A, and lysozyme. B) Representative western blot data for each protein showing that CML formation was only observed when both GO and sodium cyanoborohydride were added. MS/MS analysis showing exclusive Arg modification ([M+58]_R_, GH-DH) with GO only, but both Arg ([M+58]_R_, GH-DH) and Lys ([M+44]; [M+58]_K_, CML or GALA) modification when using GO reduction can be found in the Supporting Information. Together, these results suggest that CML formation from GO is likely an artifact of reductive amination.

Our next goal was to determine whether the [M+58] adduct observed on Lys following GO reduction corresponded to CML or a different product. To do so, we performed western blot analysis using CML-specific antibodies. Consistent with our mass spectrometry results, we found that there was a clear signal when probing with α-CML Abs only for samples that were exposed to GO reduction conditions (**Figure 4b**). These findings strongly suggest that the reported CML formation by GO has been largely artifactual, likely arising from a reduction-dependent pathway rather than direct glycation by glyoxal.

The majority of commonly used CML antibodies have been generated using carboxymethylated-KLH peptides (**Table S1**). While the exact methods used to generate CML in this case are unclear, all existing protocols we were able to find involve treatment of KLH peptides with GO and a hydride reducing agent^[42–44]^. We therefore sought to confirm that the CML adduct generated by GO reduction was the same as authentic CML generated from other glycating agents, namely glucose or ribose. To do so, we incubated peptide **1**^**K**^ with 100 mM ribose or glucose at 37 °C, which are standard in vitro conditions used to evaluate the far less reactive pentose and hexose glycating agents^[45–47]^. After 1 week, we observed modest levels of modification with ribose (17.8 ± 1.2%) but still none for glucose. To expedite our studies, we opted to accelerate this reaction by heating at 60 °C for 24 h, which produced similar levels of glycation to those observed after a week at 37 °C (**Figure S6**). We then used LC-MS to compare the extent of modification for peptide **1**^**K**^ after treatment with ribose only at 60 °C or using GO reduction (**Figure 5a**). We found that ribose produced a single [M+58] species, while peptides treated with GO reduction produced two [M+58] species, only one of which matched in retention time with that observed from ribose (**Figure 5b,c**). Given that ribose-derived [M+58] formation produces just a single Lys AGE (CML), whereas reductive amination with GO produces two distinct [M+58] products—CML and a previously reported adduct called GALA^[48]^—along with a far more abundant [M+44] adduct, we questioned whether existing CML antibodies might detect the additional [M+58] or [M+44] adducts. To evaluate this possibility, we performed western blot analysis comparing proteins glycated with GO alone, GO reduction, or ribose alone. We observed a clear signal in both the GO reduction and ribose-treated samples, confirming that authentic CML can be produced through either pathway (**Figure 5d**). Proteolytic digestion and MS/MS analysis was performed to locate sites of CML or other modifications on the protein (**Figure S5**), again showing that reduction conditions are required for CML formation with GO.

**Figure 5.**
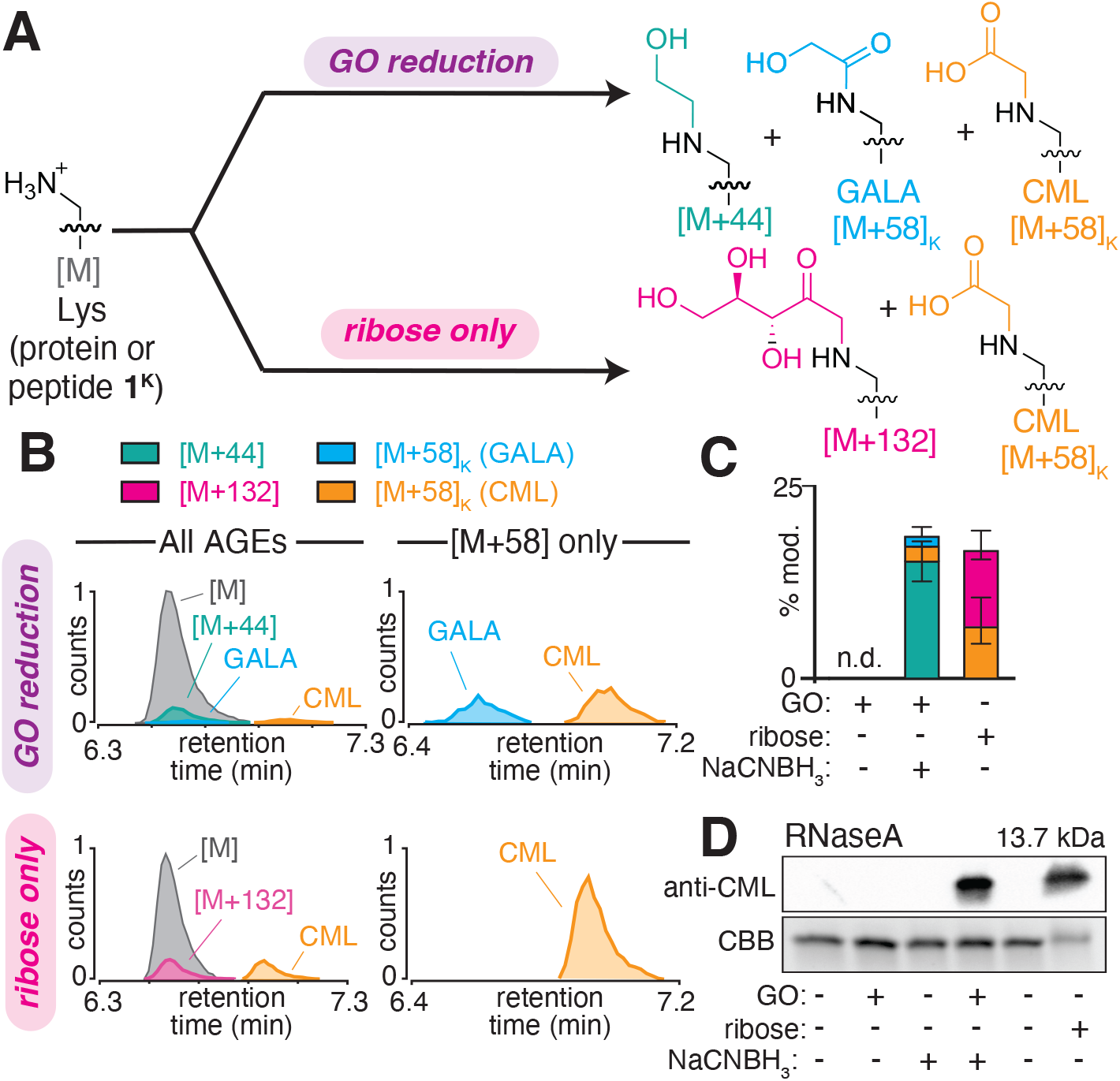
Confirming authentic CML formation with ribose. A) Reaction scheme for modification of Lys peptides (or protein substrates) by reductive amination (GO reduction), or with ribose alone. B) Representative LC-MS total compound chromatograms (TCC) for 1 mM peptide **1**^**K**^ treated with GO reduction (5 mM GO in pH 7.3 for 24 h at 37 °C, followed by a 3 h treatment with 50 mM sodium cyanoborohydride) or ribose only (100 mM ribose in pH 7.3 for 24 h at 60 °C. C) Quantified LC-MS data shows the distribution of all modified products under each conditions tested. D) Western blot analysis of RNAse A exposed to GO only, GO reduction, or ribose only shows that CML was obtained only under GO reduction or ribose only treatments.

Nonetheless, the mechanism by which reductive amination protocols generate authentic CML remains unclear. We hypothesize that hydride may facilitate CML formation by functioning as a base rather than a reducing equivalent. This model would be consistent with prior work that has suggested a deprotonation/reprotonation mechanism for glyoxalase mechanisms^[49]^. To test this possibility, we treated peptide **1**^**K**^ with 5 mM GO at pH values ranging from 7 to 12 and monitored modification over time (1-4 days.) We observed that as pH increased, [M+58] adduct formation also increased, suggesting that elevated basicity promotes CML formation (**Figure S7**). These results support the idea that CML formation under reductive amination conditions may be driven, at least in part, by increased hydride basicity rather than direct reduction by sodium cyanoborohydride.

Only two reports have contradicted the claim that GO produces CML. One study from 2004^[50]^ has been largely overlooked and has been minimally cited, though their results unambiguously align with our own. Another, from 2025^[51]^ suggests GO-derived CML formation proceeds through a hemithioacetal intermediate, but only when free thiol concentrations are well below the physiological concentrations (1–10 mM)^[52]^ of glutathione that were found to impede it. Our study is the first to definitively show that lysine modification by GO is highly dependent on the pervasive use of hydride reducing agents in past work^[11,36–39].^ Our findings strongly suggest that previously reported CML formation from GO is more accurately an artifact generated through reductive amination, rather than direct glycation under physiological conditions.

Our findings challenge conventional assumptions about GO reactivity and underscore the importance of experimental conditions in studying glycation pathways. For example, two recent metabolomics^[24]^ or proteomics^[26]^ studies have reported elevated CML levels after GO treatment, but both used hydride reducing agents. Although existing evidence strongly suggests that GO itself does not produce CML under physiological conditions, it is worth noting that alkaline conditions and/or elevated temperatures could potentially facilitate CML formation from GO in certain contexts. Additionally, the introduction of GO into a complex cellular environment may lead to interactions with other cellular metabolites, alterations in cellular localization, or modulations of pathway flux, thereby causing an increase in CML levels upon GO treatment through an indirect mechanism.

Our study redefines the landscape of GO-derived AGEs by showing not only that GO alone cannot promote appreciable levels of Lys (or Arg) carboxymethylation, but also that a single, cyclic dihydroxyimidazolidine adduct on Arg (GH-DH) is virtually the only AGE produced by GO. Although under-appreciated in past studies, GH-DH is exceptionally stable, persisting for days under physiologically relevant conditions. Additionally, it is an isomer of CMA (both [M+58]), which could easily be misattributed in proteomics analysis and entirely missed (or mis-quantified) in metabolomics studies. By redefining GO-derived AGEs and highlighting the critical role of experimental conditions in prior glycation studies, our work provides a new framework for investigating the chemical mechanisms driving protein glycation that are greatly needed to develop a complete understanding of glycation biology.

## Acknowledgments

This work was supported by National Institutes of Health grant R01GM132422 to R.A.S, as well as by a gift to the Scheck Lab from J. Kanagy, and a gift to the Scheck Lab from J. Fickel. The authors gratefully acknowledge Dr. Lynne Batchelder, Dr. Dave Wilbur, and Dr. Brian Yocis for their help with 2D NMR acquisition and analysis, and Dr. Meghan Martin along with other Scheck lab members for training on standard lab protocols.

