## Supporting Information for "A Cyclic Arginine Adduct Eclipses Carboxymethylation as the Primary Glyoxal-Derived Advanced Glycation End-Product"

Morgan E. Brutus, Colby S. Girard, Jeremiah W. Jacob-Dolan, and Rebecca A. Scheck*
Department of Chemistry, Tufts University, 62 Talbot Avenue, Medford, Massachusetts 02155

### Table of Contents

#### Supporting Materials & Methods


| | |
|---|---|
| General Materials | 3 |
| Peptide Synthesis | 3 |
| Peptide Purification | 3 |
| MALDI Mass Spectroscopy | 3 |
| Liquid Chromatography-Mass Spectrometry for Peptides | 3 |
| Liquid Chromatography-Mass Spectrometry for Intact Proteins | 4 |
| Liquid Chromatography-Mass Spectrometry for Digested Proteins | 4 |
| LC-MS Analysis and Quantification (Eq. 1) | 4 |
| General Protocol for Glycation of Peptides with Glyoxal | 5 |
| General Protocol for GO Reduction with Sodium Cyanoborohydride | 5 |
| General Protocol for Glycation of Peptides with Ribose | 5 |
| General Protocol for Protein Glycation | 5 |
| General Protocol for Dilution Experiments | 5 |
| Glyoxal Quantification by Chemical Derivatization | 6 |
| Preparation of GH-DH Modified Peptide | 6 |
| Chemical Derivatization of GH-DH | 6 |
| NMR Acquisition & Analysis | 6 |
| Western Blotting | 6 |


#### Supporting Figures and Tables


| | | |
|---|---|---|
| **Table S1** | Commercially available Carboxylmethyl Lysine (CML) monoclonal antibodies | 8 |
| **Figure S1** | Comparison of $\alpha$-oxoaldehydes, GO and MGO | 9 |
| **Figure S2** | A chemical derivatization assay to quantify 1,2 dicarbonyl concentrations | 10 |
| **Figure S3** | Chemical Derivatization of GH-DH on peptide $\mathbf{1^R}$ | 11 |
| **Figure S4** | Characterization of GH-DH on peptide $\mathbf{1^R}$ by NMR | 12 |
| **Figure S5** | AGE modifications on ubiquitin, RNAse A, and lysozyme | 13 |
| **Figure S6** | Comparison of monosaccharides, ribose and glucose | 14 |
| **Figure S7** | pH scan of GO glycation of peptide $\mathbf{1^K}$ | 15 |

### Materials and Methods

**General Materials**. All chemical reagents and solvents were of analytical grade, obtained from commercial suppliers and used without further purification unless otherwise noted. Glyoxal (GO) (40% w/v in water) (128465), Methylglyoxal (40% w/v in water) (M0252) and sodium cyanoborohydride ($NaCNBH_3$) (156159) were purchased from MilliporeSigma. D(-)-Ribose was purchased from ThermoFisher Scientific (132360250). Ribonuclease A from bovine pancreas (R6513-50MG) was purchased from MilliporeSigma. Recombinant Human HA-Ubiquitin (U-110) was purchased from R&D Systems. Lysozyme (89833) was purchased from ThermoFisher Scientific. The rabbit α-Carboxylmethyl Lysine (CML) primary antibody (CN1040) was purchased from ImmuneChem. HRP linked α-rabbit secondary antibody (7074S) was purchased from Cell Signaling Technology. Endoproteinase LysC (P8109S) was purchased from New England Biolabs. Deuterium oxide for NMR analysis was purchased from Cambridge Isotope Laboratories, Inc (DLM-4-10X0.7). All other Fmoc-protected amino acid monomers were purchased from ChemPep Inc. or Advanced ChemTech Inc.

**Peptide Synthesis.** Peptides were prepared using standard Fmoc-based solid phase peptide synthesis on Fmoc-Ala-Wang resin (100-200 mesh, 0.84 mmol/g loading, CreoSalus Inc.), typically on a 200 μmol scale in a 5 mL polypropylene fritted syringe. Subsequent amino acids were added after Fmoc-deprotection (4 mL of 20% piperidine in DMF, 2 x 15 min), and washing with DMF (4 mL, 4 x 1 min). Couplings were accomplished by incubation of 5 equiv. of amino acid, relative to resin loading, with O-(benzotriazol-1-yl)-N,N,N′,N′-tetramethyluronium hexafluorophosphate (HBTU, 5 equiv.) and DIEA (10 equiv.) for 1 hour in a total volume of 2-3 mL of DMF. These were followed by cycles of Fmoc deprotection and washing with DMF prior to the subsequent coupling step. N-terminal acetylation was achieved by incubation with a solution of acetic anhydride (1.5 equiv.) with DIEA (3 equiv.) for 2 hours in 3 mL DMF. The side chain protecting groups used were as follows: Arg(Pbf), Glu(OtBu), His(Trt), Lys(Boc), Ser(tBu), and Tyr(tBu). Side-chain acid-deprotection and peptide cleavage was conducted with trifluoroacetic acid (TFA), triisopropylsilane (TIPS) and water (95:2.5:2.5) in 5 mL of acid solution per 200 mg resin. The acid solution was then concentrated under air flow and re-dissolved in 1-3 mL of a water/acetonitrile mixture, which varied due to peptide solubility, prior to purification.

**Peptide Purification.** Peptides were purified on semi-preparative scale using an Agilent ZORBAX SB-C18 column (9.4 x 250 mm, 5 μm particle size) on an Agilent 1260 Infinity system with a water/acetonitrile mobile phase containing 0.1% TFA. Crude peptide solutions (100 μL) were injected onto the column and eluted using a standard purification gradient of 10-40% acetonitrile in water over 30 minutes at a flow rate of 3.0 mL/min. Eluting peaks were monitored by absorbance at 215 nm and 280 nm and were collected using an automated fraction collector. Collected fractions were characterized using a Bruker Microflex matrix-assisted laser desorption/ionization time-of-flight (MALDI-TOF) mass spectrometer and/or an Agilent 6530 quadrupole time of-flight (Q-TOF) mass spectrometer (see below for details) to assess peptide purity. Pure fractions were combined and lyophilized, and then each was prepared as a stock peptide solution, 20 mM in DMF.

**MALDI Mass Spectrometry.** Matrix Assisted Laser Desorption Ionization-Time of Flight (MALDI-TOF) mass spectra were obtained on a Bruker Microflex MALDI-TOF. Samples were co-crystallized using saturated solutions of α-cyano-4-hydroxycinnamic acid in 50% acetonitrile, 50% water with 0.1% trifluoroacetic acid onto ground steel plates.

**Liquid Chromatography-Mass Spectrometry for Peptides.** Reversed phase liquid chromatography and mass spectrometry (LCMS) analysis was carried out using an Agilent 1260 Infinity LC system coupled with Agilent 6530 Accurate Mass QTOF. Peptide reaction mixtures were injected onto an AdvanceBio Peptide 2.7 μm column (2.1 x 150 mm, Agilent), and eluted with a binary mobile phase of water with 0.1% formic acid (A) and acetonitrile with 0.1% formic acid (B). The elution method was as follows: isocratic 2% B from 0-1.75 min (0.400 mL/min); a gradient change of 15% B to 40% B from 1.76-12.00 min (0.400 mL/min); a isocratic change of 40% B to 100% B from 12:01-17.00 min (0.400 mL/min); and isocratic 100% B to 5% B from 17.01-26.00 min (0.400 mL/min) with column heating at 55 °C. Mass spectrometry was accomplished with an electrospray ionization (ESI) source in the positive mode and spectra in the range of 200-3000 (m/z) were collected at a rate of 5 scans/sec from 1.75-26.00 min of the chromatography method. MS acquisition was achieved using the parameters: ESI capillary voltage, 4000 V; fragmentor, 150 V; gas temperature, 325 °C; gas rate, 10 L/min; nebulizer, 40 psi.

**Liquid Chromatography-Mass Spectrometry for Intact Proteins.** Reversed-phase chromatography and mass spectrometry were performed on an Agilent 1260 Infinity LC system in line with an Agilent 6530 Q-TOF. Intact samples were diluted in water and injected onto a Zorbax 300 SB-C8 Rapid Resolution HD 1.8 μm column (2.1 x 100 mm, Agilent) and were eluted with a $H_2O$:ACN gradient mobile phase with 0.1% formic acid (0.400 mL/min; 95% - 20% water over 26 min). The mass spectrometer was utilized in positive mode with a dual electrospray ionization (ESI) source. MS spectra were acquired using the following settings: ESI capillary voltage, 4500 V; fragmentor, 250 V; gas temperature, 325°C; gas rate, 12.5 L/min; nebulizer, 50 psig. Data was acquired at rate of 5 spectra per sec and scan range of 100 – 3000 m/z.

**Liquid Chromatography-Mass Spectrometry for Digested Proteins.** Following tryptic digestion or Endoproteinase LysC digestion, peptide fragments were injected onto an AdvanceBio Peptide 2.7 μm column (2.1 x 150 mm, Agilent) and were eluted with a $H_2O$:ACN gradient mobile phase with 0.1% formic acid (0.400 mL/min; 95% - 5% water over 19 min). MS spectra were acquired using the following settings: ESI capillary voltage, 4000 V; fragmentor, 150 V; gas temperature, 325°C; gas rate, 12 L/min; nebulizer, 40 psig. Data was acquired at a rate = 5 spectra per second and scan range of 300 – 3000 m/z. MS/MS spectra were acquired using the following settings: ESI capillary voltage, 4000 V; fragmentor, 150 V; gas temperature, 325°C; gas rate 12 L/min; nebulizer, 40 psig. MS/MS was acquired at 2 spectra per second with a mass range of 100–3000 m/z, with stringency set to a medium isolation width. After identification, precursor ions were subjected to iterative rounds of collision induced dissociation in the collision chamber and subsequent mass identification. A ramped collision energy was used with a slope of 3.6 and offset of -4.8 as well as a slope of 3 and offset of 2.

**LC-MS Analysis and Quantification.** Analysis of mass spectrometry data was carried out using Agilent MassHunter Qualitative Analysis software. MS data was quantified using the MassHunter Molecular Feature Extractor, which reports cumulative ion counts (MS) as 'volumes' observed for any and all charge states associated with a particular ion. As peptides and their modified counterparts can ionize differently (eg. different charge states or different salt adducts) this method provides a more robust measure than comparing only a single charge state. For each peptide sample, compound lists were generated for each replicate of GO and ribose treatments. Quantification was carried out by dividing the AGE adduct(s) volume by the total volume of both modified and unmodified peptide (**Equation 1**). This quantification approach allows for a robust comparison of glycation extents across different AGEs on different peptide substrates, even though each may exhibit some variation in ionization efficiency.[1] Additionally, absolute counts of peptides that were not treated with GO and ribose were first evaluated to confirm that roughly similar levels of ionization were observed at the same known concentration. Retention time (RT) was used to identify discrete isomers with degenerate masses.

***Equation 1:***

$$\%\,glycation = \frac{volume\ of\ AGE\ adducts}{volume\ of\ total\ peptide}\,x\,100$$

**General Protocol for Glycation of Peptides with Glyoxal.** In general, glycation reactions for peptides in solution were performed at a 20-50 µL scale in 200 µL PCR tubes. Synthetic peptides purified in our laboratory were prepared as 20 mM stock solutions in DMF . GO stocks at this concentration were prepared by dilution of 23.0 µL of 40% w/v solution into 10 mL of ultrapure water (20 mM). To perform *in vitro* glycation on a 50 µL scale with glyoxal, to 25 µL of ultrapure water was added 10 µL of 100 mM phosphate buffered saline (PBS) at pH 7.3, 12.5 µL of a 20 mM GO stock in water, and, lastly, 2.5 µL of a 20 mM stock peptide solution in DMF. The final concentrations were 1 mM peptide and 5 mM GO in 20 mM PBS with 5% DMF co-solvent. The peptide was always added in the final step to prevent any high concentration exposure, and the glycation of each peptide was assessed individually. Tubes were capped, briefly spun in a benchtop microcentrifuge and incubated in a 37 °C water bath, typically for 24 h and in some cases for up to 1 week. After incubation, peptide samples were diluted (1:100) into 5 mM Tris Buffer pH 7.3 to quench the reaction, unless otherwise noted, and subsequently subjected to LC-MS analysis. For control experiments using methylglyoxal (MGO), the same general protocol was used, though the typical MGO concentration in glycation reactions was 1 mM instead of 5 mM.

**General Protocol for GO Reduction with Sodium Cyanoborohydride.** GO reduction procedures were performed at a ~23 µL scale in 200 µL PCR tubes. First, peptides were glycated with GO following the general protocol described above. Following this protocol, a 20 µL aliquot of the reaction mixture was transferred to a PCR tube and 3.33 µL of 300 mM $NaCNBH_3$ was added (50 mM final concentration). Tubes were capped, briefly spun in a benchtop microcentrifuge and incubated in a 37 °C water bath for 3 h. After incubation, $NaCNBH_3$ treated peptide samples were diluted (1:100) into 5 mM Tris Buffer pH 7.3 to quench the reaction and subsequently subjected to LC-MS analysis.

**General Protocol for Glycation of Peptides with Ribose.** Ribose glycation reactions were performed using similar general protocols as described above, with the following modifications: Ribose stocks were prepared by dissolving 0.1501 g of D-ribose into 1 mL of ultrapure water (1 M). To perform *in vitro* glycation on a 20 µL scale with ribose, to 13 µL of ultrapure water was added 4 µL of 100 mM phosphate buffered saline (PBS) at pH 7.3, 2 µL of a 1 M ribose stock in water, and, lastly, 1 µL of a 20 mM stock peptide solution in DMF. The resulting final concentrations were 1 mM peptide and 100 mM ribose in 20 mM PBS with 5% DMF co-solvent. Tubes were capped, briefly spun in a benchtop microcentrifuge and incubated in a 37 °C or 60 °C water bath, typically for 24 h and in some cases up to 4 days. After incubation, peptide samples were diluted (1:100) into 5 mM Tris Buffer pH 7.3 to quench the reaction, unless otherwise noted, and subsequently subjected to LC-MS analysis.

**General Protocol for Protein Glycation.** For protein reactions, lyophilized RNase A, Ubiquitin, or Lyozyme was dissolved in water to obtain a stock concentration of 250 µM. Reactions we conducted in Eppendorf tubes at a final volume of 50 µL containing 50 µM protein, 80 mM PBS pH 7.3 and 500 µM GO or 10 mM ribose at 37 °C for 24 h. Post the 24h incubation, a 20 µL aliquot was taken, and respective samples were treated with 3.33 µM of 300 mM $NaCNBH_3$ (final concentration, 50mM $NaCNBH_3$) for 3 h in a 37 °C water bath. After all reaction conditions, samples were quenched with 500 mM Tris pH 7.3 (final concentration 50 mM Tris).

**General Protocol for Dilution Experiments.** For dilution experiments described in Main Text Fig. 2, peptide $\mathbf{1}^{R}$ (1 mM) was incubated with GO (5 mM) as described above. After 24 h at 37 °C, the samples were diluted 100X in 20 mM PBS (10 µL of reaction mixture diluted in 990 µL buffer) at varying pH (6-9), and were incubated at 37 °C for up to 4 additional days, then subjected to LCMS analysis without any further dilution or purification.

**Glyoxal Quantification by Chemical Derivatization.** 10 mM stock solutions of GO and MGO were prepared as described above. Next, with those stock solutions, standard curve solutions of GO and MGO were prepared at 0 mM, 0.5 mM, 1 mM, 1.5 mM ,or 2 mM. A 40 mM stock solution of aminoguanidine HCl solution in phosphate buffer was also prepared. To a 96-well plate, 100 μL of the aminoguanidine HCl stock and 100 μL of the standard solutions were added to a well and allowed to incubate at 37°C for 5 hours. Absorbance at a wavelength of 320 nm was used to detect the aminoguanidine adduct.

**Preparation of GH-DH Modified Peptide.** GH-DH modified peptide was prepared by following "General Protocol for Glycation of Peptides with Glyoxal" as described above, but were scaled up to 500 μL total volume. The sole GH-DH modified peptide product and was purified by semi-preparative HPLC using a gradient of 0-35% acetonitrile in water over 35 min at 3.0 mL/min. Collected fractions were characterized by MALDI-TOF, pooled and lyophilized. Stocks of GH-DH modified peptide were assessed for purity by HPLC and LCMS prior to analysis by NMR.

**Chemical Derivatization of GH-DH.** To evaluate boronate ester formation for GH-DH containing peptides, 6-bromo-3-pyridinylboronic acid (PBA) was dissolved in 20 mM PBS and 50% methanol at a concentration of 200 mM. To a 25 μL aliquot of glyoxal treated peptide, was added 25 μL of the PBA stock solution (100 mM final concentration). The reaction was left to incubate for an additional 2 h at 37 °C. After incubation, samples were analyzed by Matrix Assisted Laser Desorption Ionization-Time of Flight (MALDI-TOF) mass spectrometry (Bruker Microflex MALDI-TOF). Samples were diluted 1:1 with α-cyano-4-hydroxycinnamic acid matrix (in 50% acetonitrile, 50% water with 0.1% trifluoroacetic acid), and then spotted on a ground steel plate prior to analysis.

**NMR Acquisition and Analysis.** For NMR analysis, lyophilized GH-DH modified peptide (ca. 2 mg) was dissolved in 500 μL of $D_2O$ and characterized by NMR. $^1H$ spectra were acquired with a Bruker Advance III (500 MHz, 125 MHz) spectrometer. $^1H$ chemical shifts are reported as in units of parts per million (ppm) relative to $D_2O$ (s 4.79 ppm). Data are reported as follows: chemical shift, multiplicity (s = singlet, d = doublet, t = triplet, q = quartet, m = multiplet), coupling constants (Hz), and integration).
$^1H$ NMR (500 MHz, $D_2O$) δ 8.55 (s, 1H), 7.16 (s, 1H), 7.08 (d, J = 8.1 Hz, 2H), 6.80 – 6.74 (m, 2H), 5.13 (d, 1H), 5.05 (d, 1H), 4.83 (s, 1H), 4.79 (s, 1H), 4.63 – 4.56 (m, 0H), 4.50 (dd, J = 8.8, 5.8 Hz, 1H), 4.38 – 4.29 (m, 2H), 4.22 (dq, J = 11.7, 7.1 Hz, 3H), 3.83 – 3.73 (m, 2H), 3.34 (t, J = 7.2 Hz, 1H), 3.29 (s, 0H), 3.13 (dd, J = 15.3, 6.5 Hz, 1H), 3.04 (dd, J = 15.9, 8.8 Hz, 2H), 2.85 (dd, J = 14.4, 8.7 Hz, 1H), 2.42 (h, J = 9.8 Hz, 2H), 2.09 (dd, J = 13.8, 6.4 Hz, 1H), 1.98 (s, 3H), 1.95 (s, 1H), 1.66 (s, 1H), 1.59 (s, 6H), 1.64 – 1.46 (m, 2H), 1.32 (dd, J = 7.3, 2.2 Hz, 3H), 0.85 (dd, J = 21.6, 6.3 Hz, 6H).

**Western Blotting.** At their conclusion, protein glycation reactions were treated with 6X SDS loading dye containing dithiothreitol (DTT) and boiled at 95 °C for 5 min. A precast protein gel (8–16%, miniPROTEAN TGX, Bio-Rad) was used for SDS-PAGE, and the gel was run using a Tris/glycine/SDS running buffer at 225 mV for 18 to 24 min. Transfer onto PVDF membranes was accomplished with the iBlot 2 dry blotting system (Invitrogen). After transfer, the membrane was blocked in 1× TBST (20 mM Tris, 150 mM NaCl, and 0.1% Tween) with 5% (w/v) Bovine Serum Albumin (BSA) for an hour. Primary antibody rabbit α-Carboxylmethyl Lysine (CML) (ImmuneChem, CN1040) or primary antibody mouse α-MGO (Cell BioLabs, STA-011) was added (1:1000) after blocking and incubated at 4 °C overnight with shaking. Membrane was washed three times with 1× TBST for 5 min each before adding HRP-linked α-rabbit secondary antibody (Cell Signaling Technology, 7074S) or HPR-linked α-mouse secondary antibody (Cell Signaling Technology, 7076S) diluted into 5% BSA and 1× TBST (1:2000). The membrane was incubated for 1 h with secondary antibody at room temperature before washing three times with 1× TBST for 5 min. Signal was developed using Clarity Western ECL Substrate (Bio-Rad) and imaged on a Bio-Rad ChemiDoc XRS+.

**Table S1. Commercially available Carboxylmethyl Lysine (CML) monoclonal antibodies**

| CLONE | IMMUNOGEN | VENDOR | PRODUCT# | NOTES |
|---|---|---|---|---|
| CML26 | Chemical / Small Molecule corresponding to Carboxymethyl Lysine | Abcam | ab125145 | - |
| - | Carboxymethyl lysine-KLH | ImmuneChem | CN1040 | Used in this work & Di Sanzo et. al., 2021[26] |
| NF-1G | CML-HSA | Cosmo Bio USA | KAL-KH024 | - |
| 318003 | Carboxymethyl lysine KLH | R&D Systems | MAB3247 | - |
| 6C7 | Carboxymethyl lysine KLH | Sigma-Aldrich | MABN1837 | - |

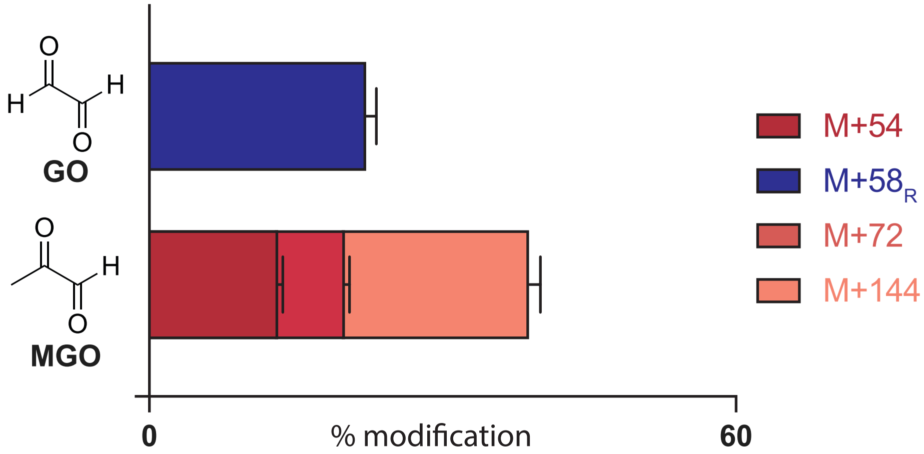


**Figure S1. Comparison of α-oxoaldehydes, GO and MGO.** Comparison of peptide **1**$^{R}$ glycation using two α-oxoaldehydes glyoxal (GO) and methylglyoxal (MGO), revealed that GO was far less reactive than MGO even when using identical reaction conditions (1 mM peptide **1**$^{R}$ treated with 1 mM glycating agent (GO or MGO) for 24 h at 37 °C). We observed 22.6 ± 0.9% total glycation when using GO but 40.1 ± 0.8% total glycation with MGO. Additionally, GO produced only a single AGE (GH-DH) while MGO produced multiple AGEs, such as MGH-1, MGH-DH, and THP.

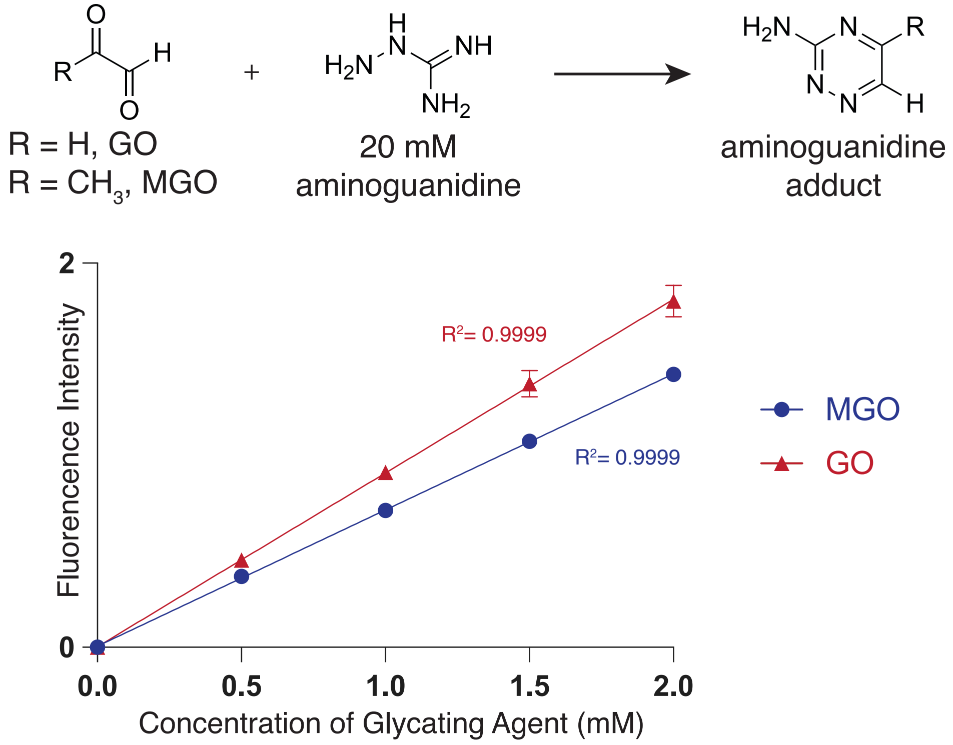


**Figure S2. A chemical derivatization assay to quantify 1,2 dicarbonyl concentrations.** Aminoguanidine binds both free MGO and GO , forming a product that can be quantified by total fluorescence intensity using a plate reader at an absorbance wavelength of 320 nm. A representative standard curve was generated using a 40 mM aminoguanidine stock solution and GO and MGO concentrations ranging from 0 to 2 mM, based on the 40% w/v in water concentration provided by the commercial vendor. The assay was completed with a high degree of linearity displayed, revealing commercial sources of GO to be 27.2 ± 0.1% higher in concentration than MGO. This suggests that GO has dampened reactivity relative to MGO.

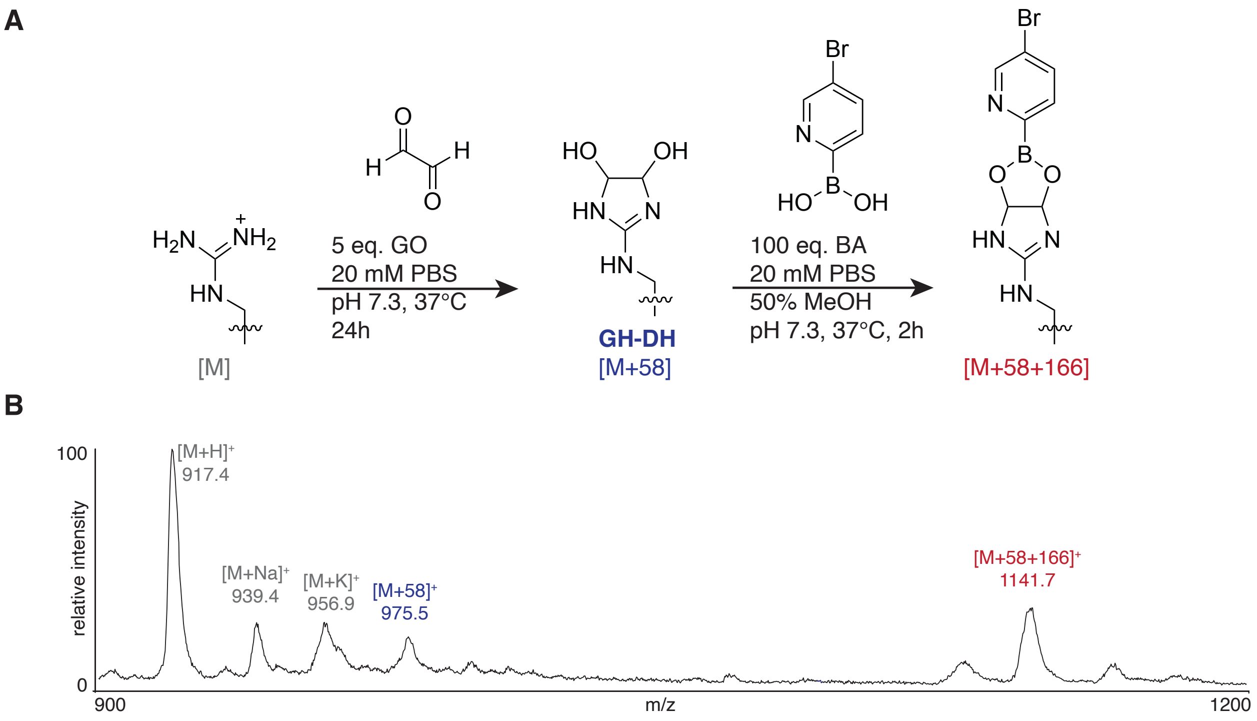


**Figure S3. Chemical Derivatization of GH-DH on peptide 1$^{R}$.** GH-DH is formed by the single addition of GO to arginine. Its structure features a vicinal diol, arising from the formation of two hemiaminals between the two aldehydes of GO and the guanidino group of Arg. (a) Scheme for the chemical derivatization of the peptide 1$^{R}$ [M+58] adduct using boronic acids. (b) Treatment of the [M+58] adduct with a brominated boronic acid resulted in formation of a new peak corresponding to the boronic ester.

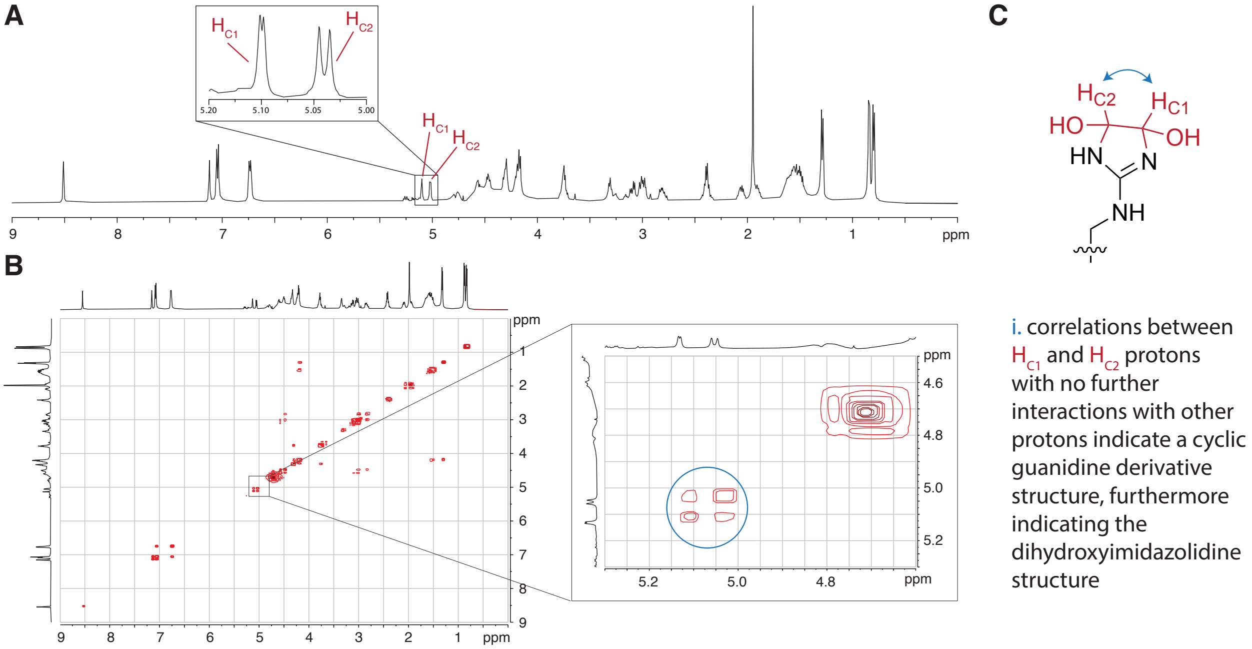


**Figure S3. Characterization of GH-DH on peptide 1^R by NMR.** Sufficient quantities of GO-modified peptide **1**$^{R}$ were prepared to allow for structural studies by NMR. (a) Using a HOD suppression experiment, a clear view of peak shifts in the region between 5.0 - 4.5 ppm was acquired. The $^{1}$H spectrum is shown with key resonances highlighted, including the GH-DH carbinol protons ($H_{C1}$ 5.13, d, 1H, $H_{C2}$ 5.05, d, 1H). These peaks show splitting patterns of doublets which would be the expected splitting of the GH-DH modification. Due to the spectra being taken in $D_2O$, the shifts of the protons on the alcohols do not appear. (b) Using COSY, it was possible to observe the correlation between the GH-DH carbinol protons ($H_{C1}$ and $H_{C2}$) which is strongly suggestive of the structure as a dihydroxyimidazolidine ring at the ωN. (c) Furthermore, there are no further cross peaks with other protons on the molecule which is also consistent with the expectation of a lack of interactions based on the dihydroxyimidazolidine structure.

ubiquitin
4 Arg
7 Lys

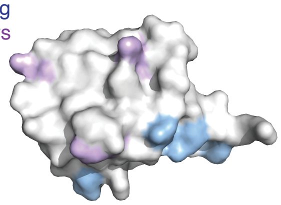

RNAse A
4 Arg
9 Lys

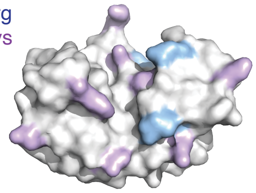

lysozyme
11 Arg
6 Lys

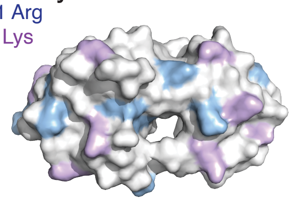

| Protein | GO only | | GO reduction | | ribose only | |
|---|---|---|---|---|---|---|
| ubiquitin | R42 | [M+58] | K6 | [M+44], [M+58] | N/A | |
| | R54 | [M+58] | K11 | [M+58] | | |
| | R72 | [M+58] | K27 | [M+44], [M+58] | | |
| | | | K48 | [M+44], [M+58] | | |
| | | | K63 | [M+58] | | |
| RNAse A | R10 | [M+58] | K1 | [M+58] | K1 | [M+58] |
| | R33 | [M+58] | K7 | [M+44], [M+58] | R10 | [M+58] |
| | R39 | [M+58] | R10 | [M+58] | K37 | [M+58] |
| | | | K31 | [M+58] | R39 | [M+58] |
| | | | R33 | [M+58] | K41 | [M+58] |
| | | | K37 | [M+58] | K61 | [M+58] |
| | | | K41 | [M+44] | K104 | [M+58] |
| | | | K61 | [M+58] | | |
| | | | K98 | [M+58] | | |
| | | | K104 | [M+58] | | |
| lysozyme | R21 | [M+58] | K1 | [M+58] | N/A | |
| | R45 | [M+58] | K13 | [M+44], [M+58] | | |
| | R68 | [M+58] | K33 | [M+44], [M+58] | | |
| | R112 | [M+58] | K96 | [M+44], [M+58] | | |
| | R125 | [M+58] | | | | |

**Figure S5. AGE modifications on ubiquitin, RNAse A, and lysozyme.** Following digestion with Lys-C, LC-MS/MS analysis was performed to determine the sites of modification for each protein, glycated either with GO alone, GO reduction conditions, or ribose alone. We found that GO only treatments resulted in exclusively Arg modification with an [M+58] mass change, corresponding to GH-DH. The GO reduction conditions substantially altered this product distribution, producing mostly, but not entirely, Lys modifications, with both [M+44] and [M+58] modifications observed. Ribose only conditions produced exclusively [M+58] adducts on both Lys (CML), and Arg.

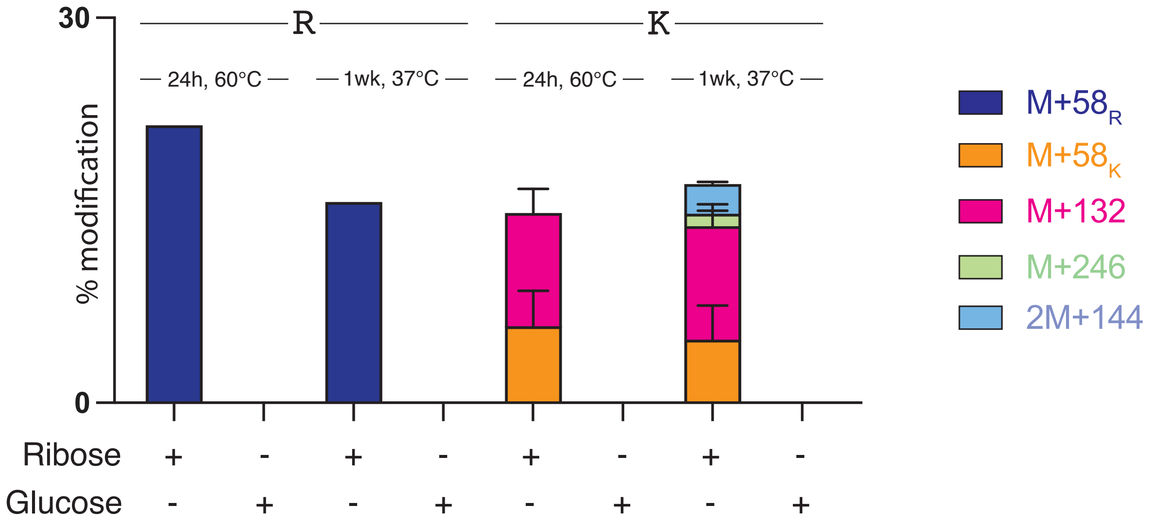


**Figure S6. Comparison of monosaccharides, ribose and glucose.** To confirm CML formation generated by the monosaccharides, ribose and glucose, we sought to see which glycating agent would allow for the formation of CML under standard glycating conditions (peptide $\mathbf{1}^{K}$ with 100 mM ribose or glucose at 37°C for 1 wk). Ribose formed AGE adducts substantially faster than glucose, which did not produce any glycation under any of the conditions tested.

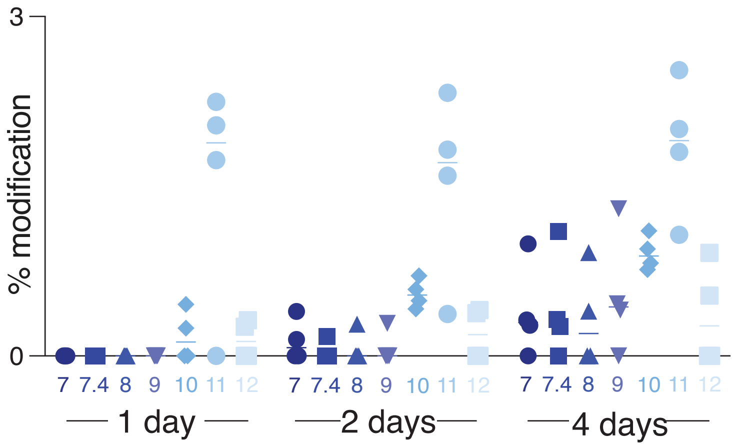


**Figure S7. pH scan of GO glycation for peptide 1$^{K}$.** Peptide **1**$^{K}$ was treated with GO at pH values from 7-12. The reaction was monitored for 1-4 days, showing increased CML formation with increased basicity, providing support for a deprotonation/reprotonation as a plausible mechanism through which GO reduction conditions can produce CML.
